# Orally administered epeleuton inhibits SARS-CoV-2 viral load, replication and pathology in the Syrian Hamster model

**DOI:** 10.1101/2021.12.07.471588

**Authors:** John Climax, Moayed Hamza, Adam Lafferty, Kate Guilfoyle, Geert van Amerongen, Konrad Stadler, Peter Wohlsein, Wolfgang Baumgärtner, Markus Weissbach, David Coughlan

## Abstract

Early treatment of patients with confirmed COVID-19 presenting mild symptoms can reduce the number that progress to more severe disease and require hospitalization. Considering the potential for the development of drug resistance to existing therapies and the emergence of new SARS-CoV-2 variants, there is a need for an expanded armamentarium of treatment options for COVID-19. Epeleuton is a novel orally administered second-generation n-3 fatty acid with potential direct antiviral and immunomodulatory actions, and a favourable clinical safety profile. In this study we show that epeleuton inhibits SARS-CoV-2 infectious viral load, replication and disease pathology in the lungs and upper airways in the Syrian hamster model of SARS-CoV-2 infection. These data support the potential utility of epeleuton in the early treatment and prevention of SARS-CoV-2 infection. Clinical trials are needed to evaluate the efficacy of epeleuton as an outpatient treatment and prevention of COVID-19.

## Introduction

Severe acute respiratory syndrome coronavirus 2 (SARS-CoV-2) is the causative agent of coronavirus disease 2019 (COVID-19) and follows SARS-CoV and MERS as the third highly pathogenic coronavirus in the last twenty years^1^. COVID-19 can range in severity from a mild respiratory illness to severe disease progressing to pneumonia, acute respiratory distress syndrome (ARDS), multi-organ failure, and death^2^.

As of early December 2021, there have been more than 262 million confirmed cases of COVID-19 globally and more than 5.2 million deaths reported to WHO. More than 7.8 billion vaccine doses have been administered. Indeed, the availability of SARS-CoV-2 vaccines has substantially mitigated COVID-19 outbreaks and aided in the global response to the COVID-19 pandemic. COVID-19 vaccination has been effective in reducing the number of new SARS-CoV-2 infections, hospitalizations and deaths^3,4^. However, uncertainties remain regarding the longevity of protection conferred by vaccination, and global disparities remain in the access and adoption of vaccines^5^. These factors contribute to a complex and varied epidemiology at both a global and regional level.

Within this evolving context, early treatment of patients with confirmed COVID-19 presenting mild symptoms can reduce the number of patients that progress to more severe disease and require hospitalization or admission to intensive care units. However, there is a paucity of therapeutic options for COVID-19 treatment in an outpatient setting and there is an urgent need for new therapeutics. Emergency use authorisations of effective therapeutics such as REGN-COV2^6^ and the emergence of molnupiravir^7^ and Paxlovid^8^ are important milestones but the need for an expanded armamentarium of therapeutic options and modalities is especially evident considering the potential for development of drug resistance to existing therapies^9^, the emergence of new SARS-CoV-2 variants^10^ and the anticipated disparities in access to newly approved treatments.

Epeleuton is 15-hydroxy eicosapentaenoic acid 15(S)-HEPE) ethyl ester, a second-generation synthetic n-3 fatty acid derivative of eicosapentaenoic acid (EPA). It is rapidly de-esterified to its active form, 15(S)-HEPE following oral administration. 15(S)-HEPE, at much lower concentrations, is an endogenous downstream 15-lipoxygenase metabolite of EPA. Epeleuton is associated with cardiometabolic and anti-inflammatory effects in clinical and preclinical studies^11^. The therapeutic effects of 15(S)-HEPE are thought to be induced by pleiotropic modes of action. As of mid-October 2021, epeleuton has completed an extensive toxicological evaluation and been administered to more than 300 subjects in phase 1 and phase 2 clinical trials, with a favourable safety and tolerability profile similar to placebo. Epeleuton is in phase 2b clinical development as a treatment for hypertriglyceridemia, type 2 diabetes and cardiovascular risk reduction (NCT04365400).

There is accumulating evidence that polyunsaturated fatty acids have anti-inflammatory, immunomodulatory and anti-viral properties, which may confer beneficial effects against SARS-CoV-2 infection^12^. Lipids play important roles in viral entry and replication, interacting with viruses throughout their life cycle. Consequently, exogenous lipid mediators may regulate the host immune response to viral infection.

Several potential binding targets for fatty acids have been identified including targets for fatty acid binding on the spike protein (S) and envelope protein of SARS-CoV-2^13, 14^. A recent study found that the S protein binds the free fatty acid linoleic acid (LA). LA binding stabilized a locked S conformation, resulting in reduced angiotensin-converting enzyme 2 (ACE2) interaction. A similar binding pocket was also present in the other highly pathogenic coronaviruses, SARS-CoV and MERS^13^. Building on this discovery it was reported that LA and EPA, the metabolic precursor of epeleuton’s active moiety 15(S)-HEPE can interfere with binding to ACE2 and significantly block the entry of SARS-CoV-2^15^.

We postulated that due to the potential of epeleuton to impact both the host immune response and to directly decrease pathogen burden by acting directly against SARS-CoV-2, epeleuton may exert a beneficial effect in the early treatment and prevention of SARS-CoV-2 infection.

In this work we determined the distribution of 15(S)-HEPE in lungs and other tissues of interest and subsequently assessed the potential of orally administered epeleuton to control SARS-CoV-2 infection and improve histopathological outcomes in the Syrian hamster model. We show that epeleuton decreases viral loads, viral RNA and pathology in both the upper and lower respiratory tract. These findings support the potential of epeleuton as an orally administered drug for the early treatment and prevention of SARS-CoV-2 infection.

## Results

### Orally administered Epeleuton accumulates in the lungs and other organs

Epeleuton (15(S)-HEPE EE) is rapidly de-esterified to its active moiety, 15(S)-HEPE following oral administration. 15(S)-HEPE is expected to exhibit similar distribution as generally known for fatty acids absorbed orally - solubilised as chylomicrons, distributed to various tissues, and released as free fatty acids through the action of tissue lipoprotein lipases^16^. Herein, we sought to investigate the tissue distribution pattern of orally administered 15(S)-HEPE EE or vehicle in Sprague Dawley rats (n=5/group) following a QDx7 dosing schedule.

Post-mortem tissue analysis (Day 8) revealed significant increases in the concentrations of 15(S)-HEPE in the tissues of treated animals compared to vehicle control (Figure 1). The mean concentrations of absorbed 15(S)-HEPE in various tissues 24 hours after the last administration of Epeleuton are as follows; Lung (6742 ng/g), heart (272 ng/g), liver (610 ng/g), kidneys (923 ng/g), skin (299 ng/g), eyes (200.96 ng/g) and spleen (1975 ng/g). On Day 8, 24 hours after the last administration, 15(S)-HEPE concentrations were below the limit of detection in the plasma and brain of treated rats in line with the plasma pharmacokinetics of the compound. While increases in 15(S)-HEPE concentrations were significant across all organs, the greatest concentrations compared to control were detected in the lungs (0ng/g vehicle control vs 6742ng/g 500mg/kg Epeleuton, P = < 0.05, Fig 1).

**Figure 1.**
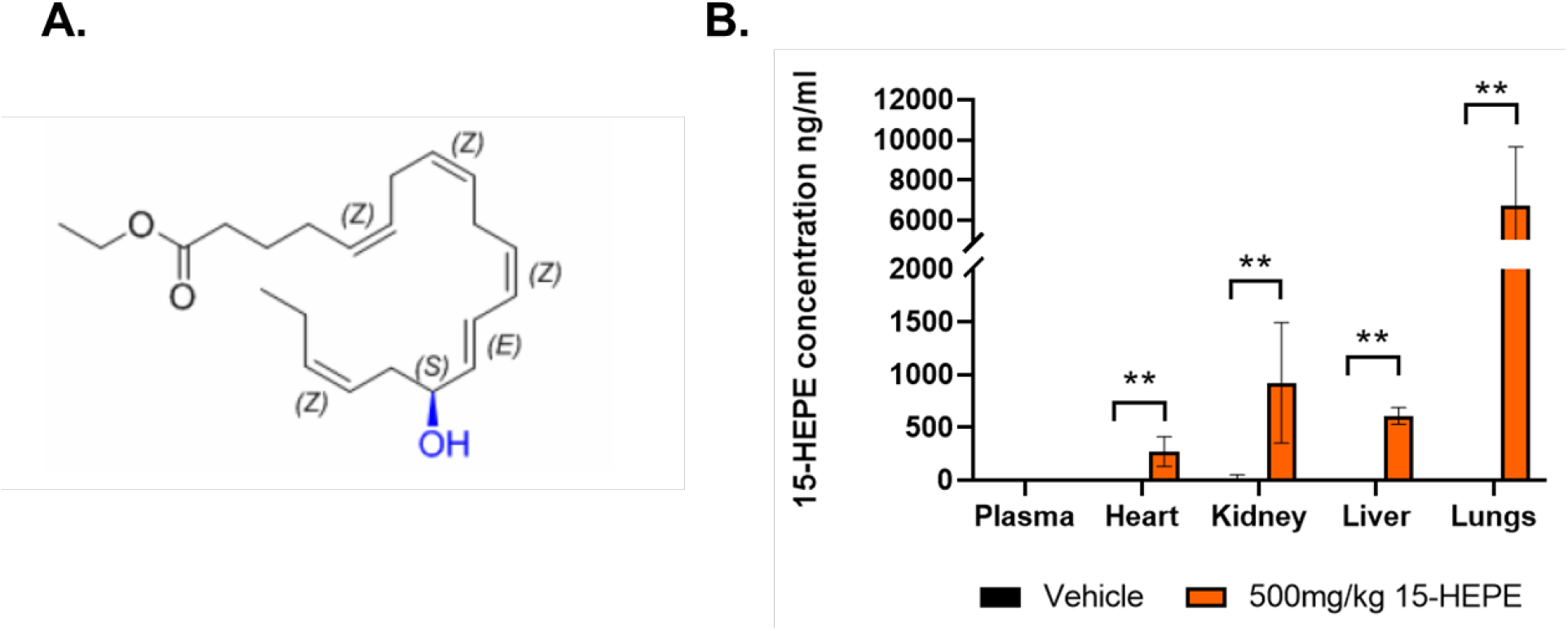
Orally administered Epeleuton accumulates in the lungs and other tissues detected as 15-HEPE. **A.** Molecular structure of epeleuton **B.** Sprague Dawley rats received 500mg/kg Epeleuton (n=5) or vehicle control (n=5) daily for 7 days. Organs were collected post-mortem, processed and subjected to LC/MS analysis to detect sequestered 15(S)-HEPE. Mean concentration is presented for each organ. Error bars represent standard deviation (S.D). Asterisk represents statistical significance versus vehicle control; * p=.033, ** p=.002.

### Epeleuton exerts a dose dependent reduction in viral replication assessed by infectious titers in hamsters challenged with SARS-CoV-2

Due to an inherent susceptibility to SARS-CoV-2 infection, Syrian hamsters are a commonly used model of SARS-CoV-2 infection kinetics and therapeutic response. However, given the self-limiting nature of SARS-Cov-2 infection in Syrian hamsters in which the virus naturally clears by day 7 post challenge (p.c) it was critical to establish steady state concentrations of epeleuton early in the course of infection, enabling assessment of its therapeutic effect and ultimately mimicking a clinical loading dose^17^.

Epeleuton dose selection was based on human equivalent doses^18^ corresponding to a 2g/day and 4g/day clinical dose, a well-tolerated clinical dosing regimen for epeleuton employed in ongoing clinical trials. All animals were administered vehicle or epeleuton (250mg/kg or 500mg/kg) once daily for 5 days prior to challenge with SARS-CoV-2. To investigate the antiviral effects of epeleuton, nine-to ten-week-old male hamsters (n=18) were inoculated intranasally with a dose of 1×10^2^ TCID_50_ SARS-CoV-2. Treatment continued for 3 days. All animals were sacrificed on day 4 p.c. for tissue collection.

Viral shedding was determined from daily throat swabs. The lower limit of quantification was determined to be 0.8 Log_10_ TCID_50_/ml, and represents undetectable viral titers. In the vehicle control arm, viral titers of 2.7 Log_10_ TCID_50_/ml were detected at day 1 (n=6). Peak viral shedding was detected on day 2 (3.6 Log_10_ TCID_50_/ml) with subsequent decreases in viral titer at days 3 and 4 (2.8 Log_10_ TCID_50_/ml and 1.8 Log_10_ TCID_50_/ml respectively, Fig. 2c).

**Figure 2.**
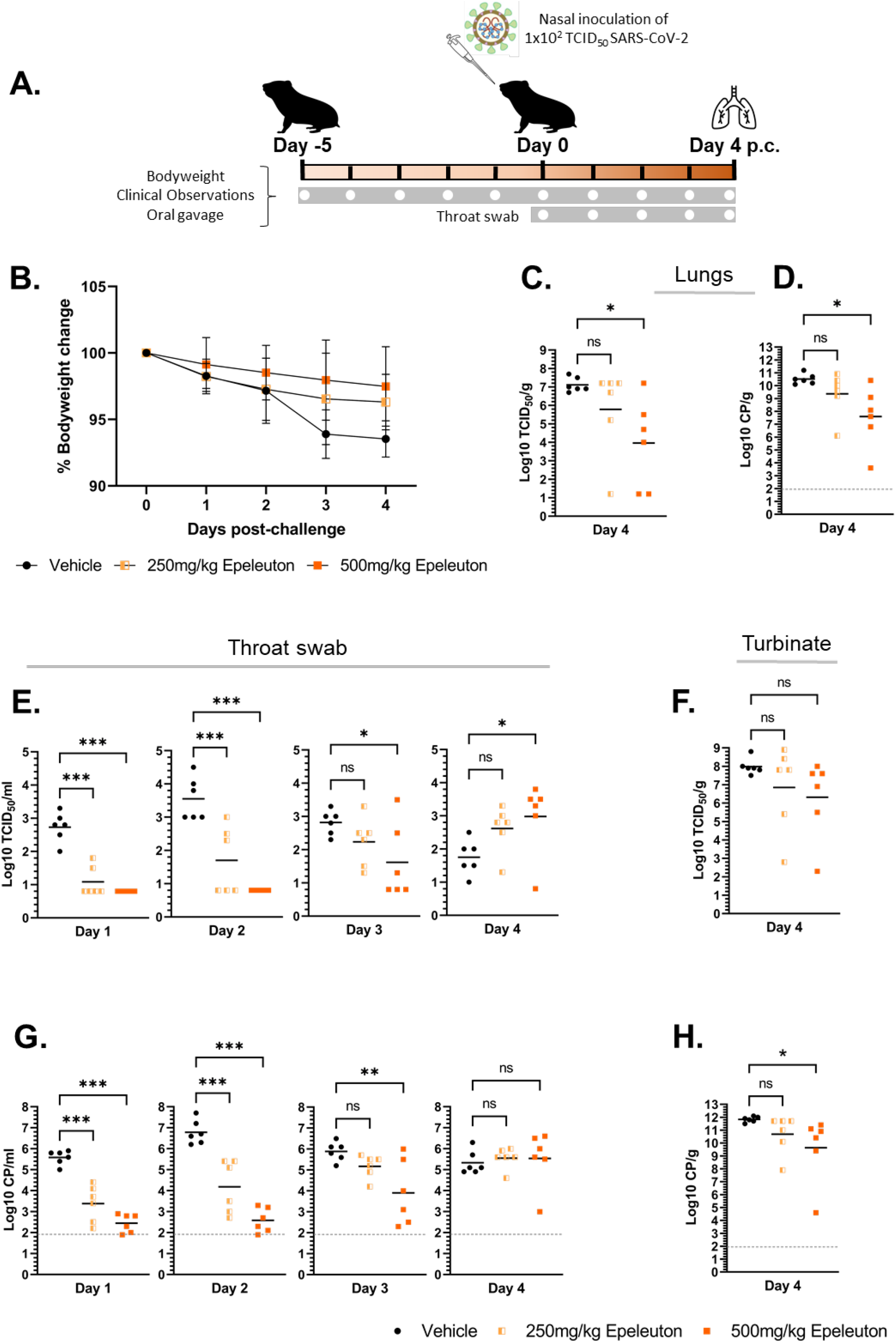
Epeleuton exerts a direct antiviral effect in a Syrian hamster model of SARS-CoV-2. **A.** Overview of the experimental design and treatment/sampling timepoints. **B.** Percentage bodyweight change in SARS-CoV-2 challenged hamsters treated with vehicle control (n=6), 250mg/kg Epeleuton (n=6) or 500mg/kg Epeleuton (n=6). **C.** Infectious viral load (Log10 TCID_50_/g) detected in lung of hamsters challenged with SARS-CoV-2 treated with vehicle control or Epeleuton collected on day 4 p.c.. **D.** Viral RNA (Log10 CP/g) detected in lung of hamsters challenged with SARS-CoV-2 treated with vehicle control or Epeleuton collected on day 4 p.c. **E.** Viral shedding assessed by viral titers (Log10 TCID_50_/ml) detected in throat swabs of hamsters challenged with SARS-CoV-2 treated with vehicle control or Epeleuton **F.** Infectious viral load (Log10 TCID_50_/g) detected in nasal turbinate tissue of hamsters challenged with SARS-CoV-2 treated with vehicle control or Epeleuton collected on day 4 p.c.. **G.** Viral shedding assessed by viral RNA (Log10 CP/ml) detected in throat swabs of hamsters challenged with SARS-CoV-2 treated with vehicle control or Epeleuton. **H.** Viral RNA (Log10 CP/g) detected in nasal turbinate tissue of hamsters challenged with SARS-CoV-2 treated with vehicle control or Epeleuton collected on day 4 p.c.. Mean concentration is presented in each graph. Error bars represent standard deviation (S.D). NS = non-significant, asterisk represents statistical significance; * p=.033, ** p=.002, *** p=<.001.

Epeleuton, when administered at a 250mg/kg dose, decreased viral shedding on days 1 and 2 characterised by significant (p = < 0.05) reductions in viral titer of 1.6 Log_10_ TCID_50_/ml and 1.9 Log_10_ TCID_50_/ml respectively. An improved effect on viral shedding was observed in animals that received 500mg/kg epeleuton. Significant decreases in viral shedding were observed up to 3 days p.c.. At both days 1 and 2, viral titers were undetectable (0.8 Log_10_ TCID_50_/ml) in animals treated with 500mg/kg Epeleuton. A significant reduction (p = < 0.05) of 1.2 Log_10_ TCID_50_/ml was observed on day 3.

Viral replication and infectious titers in the lungs and nasal turbinates were determined from tissue samples on day 4, the peak of viral load and replication. Epeleuton decreased lung viral titers in a dose-dependent manner. A mean titer of 8.0 Log_10_ TCID_50_/g was detected in the lungs of vehicle control hamsters on day 4 (Fig. 2d). Treatment with 250mg/kg epeleuton reduced viral titers in the lungs by 1.3 Log_10_ TCID_50_/g (non-significant). Epeleuton when administered at a higher dose produced a significantly improved effect reducing infectious titers in the lungs by 3.1 Log_10_ TCID_50_/g. Epeleuton produced a similar non-significant trend in nasal turbinates. Treatment with 250mg/kg and 500mg/kg epeleuton reduced viral titers in the nasal turbinates by 1.1 Log_10_ TCID_50_/g and 1.7 Log_10_ TCID_50_/g respectively (Fig. 2E).

### Epeleuton exerts a dose dependent reduction in viral replication assessed by viral RNA in hamsters challenged with SARS-CoV-2

As a secondary measure of viral shedding, viral RNA genome copy number was quantified in daily throat swabs, nasal turbinates and lung tissue. The lower limit of quantification was determined to be 1.9 Log10 copies/ml/g, and represents undetectable viral RNA. In the vehicle control arm, viral RNA of 5.6 Log_10_ copies/ml were detected at day 1 (n=6). Peak viral RNA was detected on day 2 (6.8 Log_10_ copies/ml) with subsequent decreases in viral RNA at days 3 and 4 (5.9 Log_10_ copies/ml and 5.3 Log_10_ copies/ml respectively, Fig. 2F).

Epeleuton administered at doses of 250mg/kg and 500mg/kg resulted in significant dose dependent reductions (p = < 0.05) in viral RNA up to 3 days p.c (500mg/kg). Epeleuton, when administered at a 250mg/kg dose, exerted an antiviral effect on days 1 and 2 characterised by significant (p=<0.05) reductions in viral RNA of 2.2 Log_10_ copies/ml and 2.6 Log_10_ copies/ml respectively. An improved effect on viral shedding was observed in animals that received 500mg/kg epeleuton. Treatment significantly reduced viral shedding (measured by viral RNA) on days 1, 2 and 3 by 3.1 Log_10_ copies/ml, 4.2 Log_10_ copies/ml and 2.0 Log_10_ copies/ml respectively.

Viral RNA in the lungs and nasal turbinate was determined at day 4 p.c.. A mean titer of 10.5 Log_10_ CP/g was detected in the lungs of vehicle control hamsters. Treatment with 250mg/kg epeleuton reduced viral RNA in the lungs by 1.1 Log_10_ CP/g (non-significant). This effect was seen to be dose dependent (Fig. 2G). When administered at a higher dose, epeleuton produced a significantly improved effect (p = < 0.05) reducing viral RNA in the lungs by 2.9 Log_10_ CP/g. Epeleuton produced a similar dose-dependent trend in nasal turbinates. Treatment with 250mg/kg reduced viral RNA in the nasal turbinates by 1.1 Log_10_ CP/g (non-significant). An improved significant response (p = < 0.05) was observed following treatment with 500mg/kg epeleuton. Viral RNA was significantly reduced by 2.2 Log_10_ CP/g compared to vehicle (Fig. 2H).

### Epeleuton ameliorates upper and lower airway SARS-CoV-2 pathology and inflammation

Challenge with SARS-CoV-2 resulted in a mean weight loss of 6% in the vehicle arm. Epeleuton attenuated SARS-CoV-2 related weight loss in a dose dependent manner (Fig. 2b). Consistent with the known disease severity in SARS-CoV-2 infected hamsters no other clinical signs were observed in any treatment group. All animals (n = 18) survived the viral challenge until the scheduled time point of autopsy.

Upper and lower respiratory tract involvement is the most common manifestation of SARS-CoV-2 infection, resulting in significant morbidity and mortality^19^. In order to assess the effects of epeleuton against the outcomes of SARS-CoV-2 infection both nasal turbinate and lung tissue were harvested and processed for histopathological analysis at day 4 p.c., the peak of lung viral load and replication in the Syrian hamster model. A summary of histological scores is presented in table S3. No lungs showed definite evidence of interfering concurrent infections (e.g., bacteria or fungi), nor was aspiration of foreign material clearly observed.

All animals except one (500mg/kg Epeleuton) showed inflammatory changes in the nasal cavity (Fig. 3a) consisting of infiltrating heterophilic granulocytes, macrophages and lymphocytes. The most severe rhinitis was present in the vehicle arm (score: 2.83) with decreased rhinitis observed in the 250mg/kg (score 2.0), and 500mg/kg (score 1.17) epeleuton groups. Overall, the severity of rhinitis was significantly (p=<0.05) reduced in animals treated with the higher dose of epeleuton compared to vehicle control.

**Figure 3.**
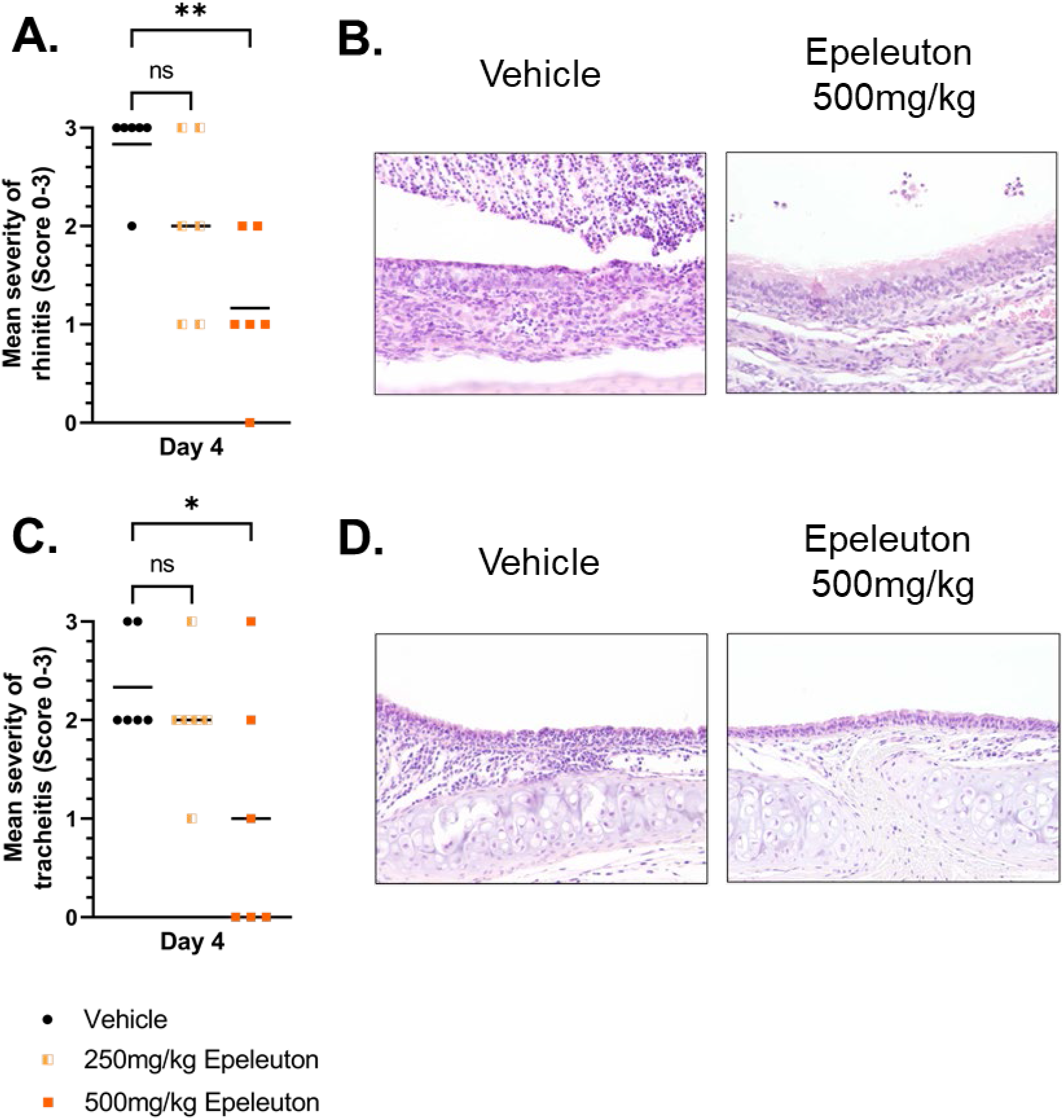
Epeleuton reduces inflammation and histopathological changes in the upper airways. **A.** Histopathological scoring of the severity of rhinitis in tissue of hamsters challenged with SARS-CoV-2 treated with vehicle control (n=6) or Epeleuton (n=6/arm). **B.** H&E Images are representative sections taken from n=1 animal/group at 200x **C.** Histopathological scoring of the severity of tracheitis in tissue of hamsters challenged with SARS-CoV-2 treated with vehicle control (n=6) or Epeleuton (n=6/arm). **D.** H&E Images are representative sections taken from n=1 animal/group at 200x. Rhinitis / tracheitis severity: 0 = no inflammatory cells, 1 = few inflammatory cells, 2 = moderate number of inflammatory cells, 3 = many inflammatory cells. Mean is presented in each graph. NS = non-significant, asterisk represents statistical significance; * p=.033, ** p=.002.

All animals, except three in the 500mg/kg epeleuton treated arm, showed inflammatory changes of the trachea (Fig. 3c) consisting of infiltrating heterophilic granulocytes, macrophages and lymphocytes. The severity of tracheitis was highest in the vehicle group (score: 2.33) with decreased tracheitis severity observed in both the 250mg/kg epeleuton (score: 2.0), and 500mg/kg epeleuton (score: 1.0) group. The severity of tracheitis was significantly (p=<0.05) reduced in animals treated with the higher dose of epeleuton compared to vehicle control.

Autopsy studies of patients who have died from severe SARS CoV-2 infection reveal presence of alveolar wall injury and diffuse alveolar damage consistent with acute respiratory distress syndrome (ARDS)^20, 21^. Most animals showed inflammatory changes in the pulmonary parenchyma (Figs. 4a-4g). Alveolitis was characterized by infiltration of heterophilic granulocytes and macrophages. The most severe alveolitis (Fig. 4b) was seen in the vehicle arm (mean: 1.33) with decreased alveolitis severity observed in the 250mg/kg epeleuton and 500mg/kg epeleuton groups. A similar pattern is observed when summarizing the scores for severity and extent of alveolitis (Fig. 4a): vehicle control (mean score: 2.16), 250mg/kg epeleuton (mean score: 0.83), and 500mg/kg epeleuton (mean score: 0.34). Reductions in both extent (−80%) and severity (−87%) of alveolitis were significant for the high dose epeleuton group only.

**Figure 4.**
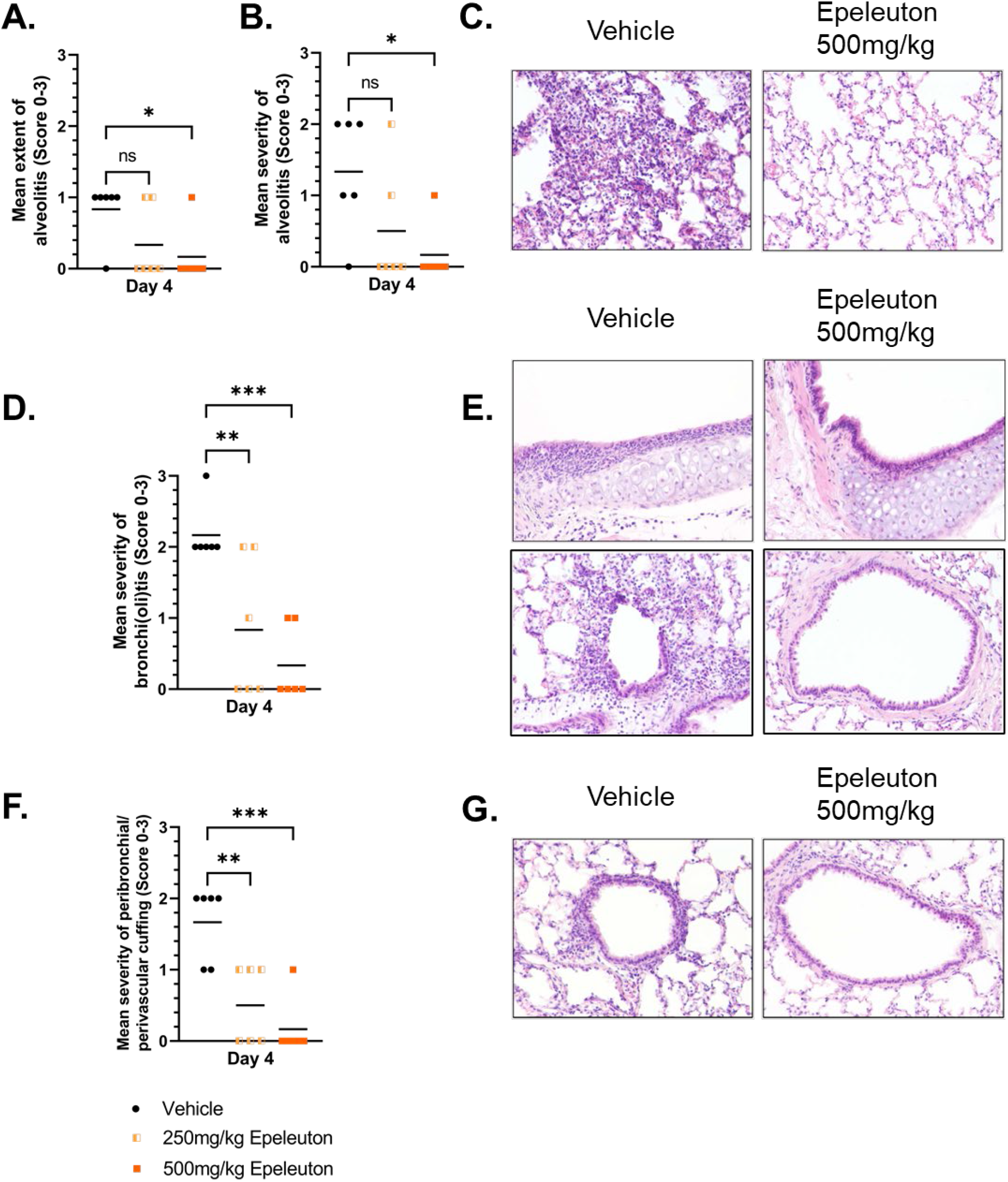
Epeleuton reduces inflammation and histopathological changes in the lower respiratory tract. Histopathological scoring of the extent of alveolitis (**A.**) and severity of alveolitis (**B.**) in tissue of hamsters challenged with SARS-CoV-2 treated with vehicle control (n=6) or Epeleuton (n=6/arm). **C.** H&E Images are representative sections taken from n=1 animal/group at 200x **D.** Mean severity of bronchitis/bronchiolitis in tissue of hamsters challenged with SARS-CoV-2 treated with vehicle control (n=6) or Epeleuton (n=6/arm). **E.** H&E Images are representative sections sections of bronchi and bronchioles taken from n=1 animal/group at 200x **F.** Percentage of sections positive for peribronchial/perivascular cuffing in tissue of hamsters challenged with SARS-CoV-2 treated with vehicle control (n=6) or Epeleuton (n=6/arm). **G.** H&E Images are representative sections taken from n=1 animal/group at 200x Alveolitis / bronchitis / bronchiolitis severity: 0 = no inflammatory cells, 1 = few inflammatory cells, 2 = moderate number of inflammatory cells, 3 = many inflammatory cells. Alveolitis extent: 0 = 0%, 1 = < 25%, 2 = 25 50%, 3 = > 50%. Extent of peribronchial /perivascular cuffing: 0 = none, 1 = 1 2 cells thick, 2 = 3 10 cells thick, 3 = > 10 cells thick. Mean is presented in each graph. NS = non-significant, asterisk represents statistical significance; * p=.033, ** p=.002, *** p=<.001.

Additional inflammatory lesions consisted of rather mild changes of bronchioles but more severe inflammatory changes of the large bronchi (Fig 4d). Besides single cell necrosis with desquamation into the lumen infiltrations of heterophilic granulocytes and macrophages were present. The most severe inflammation of the lower airways was observed in the vehicle arm (mean score: 2.17) with decreased lower airway inflammation observed in the 250mg/kg epeleuton (mean score: 0.83) and 500mg/kg epeleuton (mean score: 0.33) arms. The dose dependent reduction in severity of inflammation in the lower airways was significant for both doses of epeleuton. Perivascular and peribronchiolar cuffing with infiltration of lymphocytes, macrophages, and less heterophils occurred in all groups to a variable extent (Fig. 4f). The most severe changes were observed in the vehicle arm (mean score: 1.67) followed by a significant (p = < 0.05) dose dependent reduction in both 250mg/kg epeleuton (mean score: 0.50) and 500mg/kg epeleuton (mean score: 0.17) groups.

## Discussion

In this study, we used the Syrian hamster model of SARS-CoV-2 infection to assess the effect of epeleuton, a second-generation novel n-3 fatty acid derivative of EPA on SARS-CoV-2 infectious viral load, replication and pathology. Epeleuton substantially reduced the replication of SARS-CoV-2 in the lungs and upper airways based on both viral RNA genome copy number and infectious virus titers. Notably, at day 4, the peak of lung viral load in the Syrian hamster model there was significant inhibition of viral replication in the lung based on both infectious titers and viral RNA. This corresponded with markedly reduced lung and upper airway pathology, indicating improved disease outcome and potential for complete respiratory tract protection. Importantly, the observed decrease in extent and severity of alveolitis and lower airway inflammation may represent an important finding indicating therapeutic potential considering that one third of patients hospitalised with COVID-19 meet ARDS criteria and that autopsies from patients who died following severe COVID-19 reveal alveolar wall injury and diffuse alveolar damage^19, 20, 21^.

SARS-CoV-2 and other coronavirus’ pathology like all RNA viruses is complex and impacted by both host and pathogen burden. Epeleuton as a polyunsaturated fatty acid is expected to employ pleiotropic modes of action to influence SARS-CoV-2 viral entry, viral replication and disease outcome. A recent review by Theken et al. outlined the important roles that lipids play in the viral life cycle both through direct interactions with viruses and by modulating various aspects of host biology^12^. As such, the observed effects of epeleuton in the Syrian hamster model are likely to consist of both direct antiviral effects and protection conferred by epeleuton’s effects on host immune response, including its known anti-inflammatory effects^11^. Recent studies suggest several potential targets for fatty acid binding to SARS-CoV-2 and provide evidence that polyunsaturated fatty acids can exert direct antiviral effects independent of host factors. The identified binding targets include a free fatty acid (FFA) binding pocket on the spike protein (S)^13^ and a potentially druggable target on the transmembrane domain of the envelope protein (E) of the SARS-CoV-2 virus^14^. Toelzer et al. found that linoleic acid (LA), an omega-6 fatty acid was tightly bound to the newly discovered FFA binding pocket. LA binding stabilized a locked conformation of the S protein, resulting in reduced ACE2 interaction. A similar binding pocket was also present in the other highly pathogenic coronaviruses SARS-CoV and MERS^13^. Similarly, a fatty acid binding pocket was previously exploited to develop a small-molecule antiviral therapeutic to treat rhinovirus which was successful in human clinical trials^22^. It is noteworthy that addition of LA and arachidonic acid, another polyunsaturated fatty acid, decreased replication of both MERS-CoV and HCoV-299E in *in vitro* experiments^23^.

Building on the discovery of the FFA binding pocket, using a spike protein pseudo-virus, it was demonstrated that both omega-3 and omega-6 polyunsaturated fatty acids including LA and EPA interfere with binding to ACE2 and significantly block the entry of SARS-CoV-2^15^. EPA is the metabolic precursor of epeleuton’s active moiety 15(S)-HEPE. Importantly, the presence of a conserved FFA binding pocket among various pathogenic coronaviruses suggests that this druggable target may be conserved in emerging SARS-CoV-2 variants, and offers hope that if proven effective compounds targeting the FFA binding pocket may be less susceptible to the development of drug resistance.

The observed inhibitory effects of epeleuton against SARS-CoV-2 viral replication may also be explained in part by its effect on host biological pathways and immune response. In fact, the importance of metabolic pathways especially lipid metabolism on viral entry into cells may also partly explain the higher risk of severe disease among individuals with obesity and pre-existing cardiovascular disease^24^. In a phase 2a clinical trial of epeleuton in patients with obesity and NAFLD, serum biomarker data provides several hints regarding systemic effects of epeleuton on the host lipidome which may impact SARS-CoV-2 replication. In this phase 2a clinical trial epeleuton significantly decreased concentrations of plasma free fatty acids^11^, an observation that may be partly explained by the known inhibition of fatty acid synthase (FAS) by polyunsaturated fatty acids^25^. FAS is a homodimeric multifunctional enzyme that catalyzes fatty acid synthesis, synthesising palmitate from acetyl-CoA and malonyl-CoA in the presence of NADPH^26^. In turn, palmitoylation modification of the SARS-CoV-2 spike protein has been shown to be an important step for viral infectivity, allowing the virus to interact with cell membranes and initiate viral fusion^27^. In a recent study Chu et al. demonstrated that FAS inhibitors could reduce cellular levels of free fatty acids and palmitoylated proteins. They subsequently demonstrated that FAS inhibitors can inhibit SARS-CoV-2 replication in human cell lines and hACE2 transgenic mice, and consequently that cellular lipid synthesis is required for SARS-CoV-2 replication^28^.

Although future investigations will need to establish the relative contributions and mechanisms of direct antiviral effects and alterations of host biological pathways to epeleuton’s efficacy against SARS-CoV-2, the outcomes observed in the present model, the clinical anti-inflammatory effects of epeleuton and the extensive literature supporting opportunities for therapeutic intervention against SARS-CoV-2 with polyunsaturated fatty acids are encouraging. Notably, epeleuton is furthest in clinical development as a cardiometabolic treatment (NCT04365400) and given the contributions of vascular and endothelial dysfunction to the persistent and prolonged effects of COVID-19^29,30^, epeleuton may hold additional promise in the prevention and amelioration of long COVID.

A limitation of the present study is that treatment was initiated pre-infection to account for potential pharmacokinetic differences between hamsters and other species used in previous studies of epeleuton, and to ensure steady state levels are achieved early in the course of infection. Polyunsaturated fatty acids may take days to reach therapeutic concentrations in plasma and desired tissues. Consequently, while the observed results indicate that epeleuton exerts protective effects against SARS-CoV-2 infection and end-organ outcomes, the study design leaves some uncertainty regarding the potential treatment window. However, the marked effects of epeleuton on lung and upper airway pathology are consistent with previous findings that direct acting antivirals including for SARS-CoV-2 tend to improve disease outcome when administered early in the course of infection^31, 32^.

To overcome the pharmacokinetic challenge posed by the acute course of SARS-CoV-2 infection, Cardiolink-9, a recent small exploratory phase 2a clinical trial of icosapent ethyl, an ethyl ester formulation of EPA, in patients with COVID-19 employed a loading dose. This trial provided the first human experience with a 4g BID (8 g/day) loading dose of a fatty acid therapy. The trialled treatment regimen demonstrated favourable short-term safety and tolerability, comparable to usual care, and resulted in significant improvements in inflammatory markers and patient symptom scores from baseline^17^. Since EPA is the metabolic precursor of 15(S)-HEPE this trial provided a template dosing strategy for epeleuton. Future clinical trials could employ a similar strategy with a loading dose of epeleuton to rapidly achieve therapeutic concentrations of 15(S)-HEPE.

Although epeleuton decreased both upper airway and lung pathology while simultaneously substantially decreasing viral replication in the lungs and attenuating body weight loss, an apparent increase in viral shedding is observed in the throats of hamsters at day 4 after infection. The significance of this finding seems tenuous because importantly it contrasts to the marked and consistent improvements in disease outcome across both the upper and lower respiratory tract. Further characterisation of the antiviral mechanism of action of epeleuton against SARS-CoV-2 may aid in ascertaining the relevance of this observation. However, previous studies in the Syrian hamster model indicate that the effects of treatment on viral replication in the lung are more relevant than viral shedding to disease outcome. The nucleoside analog MK-4482’s (molnupiravir) hamster and clinical data provide supportive evidence. In the Syrian hamster model, molnupiravir inhibited both lung pathology and viral replication in the lung but did not decrease viral shedding measured with oral swabs at days 2 and 4^32^. Encouragingly molnupiravir’s efficacy in the hamster model was replicated clinically, with results indicating that early treatment with molnupiravir reduced the risk of hospitalization or death compared to placebo for patients with mild or moderate COVID-19^7^.

The need for an expanded armamentarium for the treatment of COVID-19 is especially evident considering the potential for the development of drug resistance to existing therapies and the emergence of new SARS-CoV-2 variants^9^. Combination therapy utilising drugs with different mechanisms of action may add to the treatment options available and may be more effective than single agents by exploiting potentially additive effects. Instructive precedents include combination approaches for the treatment of human immunodeficiency virus and hepatitis C virus which yielded improved efficacy compared to single agents as well as promising combination approaches for treating MERS-CoV^33, 34, 35.^ Although the mechanism of action of epeleuton against SARS-CoV-2 requires further characterisation, it is anticipated that epeleuton acts through a distinct and unique mechanism compared to currently available therapeutics such as the nucleoside analogs remdesivir and molnupiravir, the REGN-COV2 monoclonal antibody combination, Paxlovid and corticosteroids. It is noteworthy that following the discovery of the FFA binding pocket on the S protein, *in vitro* experiments revealed that supplementation of the polyunsaturated fatty acid LA synergizes with remdesivir, suppressing SARS-CoV-2 replication more effectively than with remdesivir alone^13^.

Epeleuton is orally administered with a favourable safety and tolerability profile^11^. Epeleuton’s oral route of administration and stability at room temperature provide the possibility of early treatment and the potential for distribution to both high and low resource regions of the world in the fight to reduce COVID 19 disease burden. Currently, there is a paucity of oral therapeutic options for COVID-19 treatment in an outpatient setting and there is an urgent need for new therapeutics. The present findings support the potential of epeleuton as a novel orally administered drug for the early treatment and prevention of SARS-CoV-2 infection. Further studies will be required to elucidate the therapeutic and preventative potential of epeleuton against SARS-CoV-2 infection.

## Acknowledgements

The authors would like to acknowledge the laboratory, animal technicians and wider team at Viroclinics Xplore for their work on the hamster efficacy study.

## Methods

### Epeleuton and dose vehicle

15(S) HEPE-EE (15-Hydroxy-eicosapentaenoic acid ethyl ester) was supplied by Afimmune Limited.

Hydroxypropyl methyl cellulose (HPMC) (Methocel E4M) was supplied by IMCD UK Ltd. The dose vehicle was prepared as a 0.5% (w/v) aqueous solution and stored in a refrigerator set to maintain a temperature of +2-8°C until use.

### 15-HEPE Tissue distribution Study

#### Compliance with Ethical Standards

Ten male Sprague Dawley rats were used in this study. Animals were supplied by Charles River Ltd., Margate UK. Animals were acclimatised to the experimental unit for at least 5 days prior to study commencement. During the pre-trial and on-study periods, animals were group housed in caging appropriate to the species. A standard laboratory diet (SDS RM1 € SQC, batch 2193) and mains quality tap water were available *ad libitum* throughout the study.

#### Tissue distribution study protocol

Eleven male Sprague Dawley rats (n=5/group) received a once daily oral dose of 500mg/kg Epeleuton or vehicle for 7 days. Animals were observed 15, 30 and 60 minutes following each dose administration. On day 8, 24 hours following the final daily dose (on day 7), animals were humanely sacrificed and samples collected.

#### Sampling

Approximately 24hrs following completion of the dosing regimen, on Day 8, a terminal blood sample was collected from each animal from the orbital sinus (under anaesthetic). The animals were then humanely sacrificed by CO_2_ narcosis and the following tissues removed, weighed, snap frozen in liquid nitrogen and stored in a freezer set to maintain a temperature of −80°C until analysis:

Lungs (perfused with phosphate buffer saline), heart, brain, eyes, liver, kidney, spleen, colon and skin (shaved).

Blood samples were centrifuged (1500g, 10min, +4°C) and the resultant plasma stored in a freezer set to maintain −80°C until analysis.

##### Plasma sample preparation and analysis

For each batch, calibration samples were freshly prepared in 5% (w/v) Bovine Serum Albumin (BSA) in phosphate buffered saline (PBS) to prevent the potential effect of the 15(S)-HEPE endogenous levels on the assay.

The standards, quality control and test samples were prepared, extracted and analysed in batches using research grade methodology detailed in the following sections.

Test samples resulting in a determined concentration below the lowest calibration standard were reported as less than the lower limit of quantification (<LLOQ).

##### Plasma extraction procedure

15(S)-HEPE and its internal standard 15(S)-HETE-d_8_ were determined from rat plasma by enzymatic digestion followed by protein precipitation using validated bioanalytical method.

##### Tissue extraction procedure

15(S)-HEPE and its internal standard 15(S)-HETE-d_8_ were determined from rat tissues by enzymatic digestion followed by protein precipitation using validated bioanalytical method.

#### Key analytical equipment

Mass spectrometer: API4000, Sciex.

HPLC system: PE 200 pumps, Perkin Elmer

Autosampler: HTC PAL,

Data handling system: Analyst Version 1.6.2, Sciex.

Laboratory Information Management System: Watson 7.4.2, Thermo Scientific.

Analytical column: Poroshell EC-C18, 50 x 2.1mm, 2.7 μm,

#### Key mass spectrometer parameters

Ionisation Mode: TurboIonSpray

Q1 Resolution: Unit

Q3 Resolution: Unit

Ions Monitored:

| Compound | Q1 (m/z) | Q3 (m/z) | Polarity | Dwell Time (msec) |
| --- | --- | --- | --- | --- |
| 15(S) HEPE | 317.3 | 219.2 | Positive | 250 |
| 15(S) HETE-d<sub>8</sub> | 327.4 | 226.3 | Positive | 250 |

#### Key chromatographic parameters

Mobile Phase: Acetonitrile/water/acetic acid (50/50/0.5, v/v/v)

***Key chromatographic parameters*** Mobile Phase: Acetonitrile/water/acetic acid (50/50/0.5, v/v/v)
| Time (mins) | %A | Flow Rate (µL/min) |
| --- | --- | --- |
| 0.0 | 50 | 750 |
| 2.5 | 50 | 750 |

#### Data calculations

Data manipulations carried out during this phase of the study were performed using Sciex Analyst Version 1.6.2 data handling software, combined with Thermo Scientific Watson™ Version 7.4.2 LIMS processing and reporting software. 15(S) HEPE and internal standard peak area and retention time data were obtained using Analyst. Calibration and quantification resulting in the calibration parameters (slope, intercept and coefficient of determination) and reported calibration standard, quality control and test sample determined concentrations were performed within Watson™ using the 15(S) HEPE and internal standard peak area data imported electronically from Analyst. Calibration standard and quality control sample matrix concentrations were entered into Watson™ to 3 significant figures. 15(S) HEPE determined concentrations relating to test samples are reported to 3 significant figures.

### *In vivo* efficacy study

#### Animals

Male Syrian golden hamsters, nine to ten weeks old, were obtained from Janvier Labs (France). They were housed and all procedures were performed at the Viroclinics Xplore animal facility (Schaijk, The Netherlands), under conditions that meet the standard of Dutch law for animal experimentation and in accordance with Directive 2010/63/EU of the European Parliament and of the Council of 22 September 2010 on the protection of animals used for scientific purposes. Ethical approval for the experiment was registered under protocol number: AVD27700202114492-WP04.

All procedures were performed under sedation with isoflurane (3-4%/O2) and animals were weighed and observed daily for changes in physical appearance or behaviour.

#### Treatment

On day −5 to day −1, and days 0 to 3 p.c., animals were treated under anaesthesia once daily via the oral route of administration (p.o.), with a dose of 250 or 500mg/kg body weight. Animals were weighed daily prior to treatment and the dose volume was adjusted accordingly.

#### Challenge

On day 0, all animals were infected via the intranasal route of administration (i.n.) with 10^2.0 TCID50 SARS-CoV-2 (BetaCoV/Munich/BavPat1/2020) in a total dose volume of 100μl, divided equally over both nostrils. Animals were followed for four days and on day 4 p.c., were euthanised by abdominal exsanguination under anaesthesia.

#### Sampling of the respiratory tract

Throat swabs were collected prior to infection on day 0 and then daily during the infection phase, days 1 to 4 p.c.. Upon necropsy on day 4 p.c., lung and nasal turbinate tissue samples were collected. Left lung lobes and nasal turbinates and trachea were preserved in 10% neutral buffered formalin for histopathology. The right lung lobes and nasal turbinate were homogenised, briefly centrifuged, aliquoted and frozen for subsequent virological analyses.

#### Detection of replication competent virus

Quadruplicate, 10-fold serial dilutions of throat swabs and tissue homogenates were transferred to 96 well plates with confluent layers of Vero E6 cells and incubated for one hour at 37°C. Cell monolayers were washed and incubated for five days at 37°C. Plates were then scored using the vitality marker WST8 and viral titers (log10 TCID50/ml or /g) were calculated using the method of Spearman-Karber.

#### Detection of viral RNA

Viral RNA was isolated from the throat swabs and tissue homogenate samples and Taqman PCR was performed using specific primers and probe for beta coronavirus E gene. The number of copies (Log10 CP/ml/g) in each sample was calculated against a standard included in each of the qPCR runs performed.

#### Gross pathology

At the time of necropsy, all lung lobes were inspected and an estimation of the percentage affected lung tissue from the dorsal view was described.

#### Histopathology

Histopathological analysis was performed on selected tissues (lung, trachea and nasal turbinates). After fixation with 10% formalin, sections were embedded in paraffin and the tissue sections were stained with haematoxylin and eosin (H&E) stain for histological examination. Histopathological assessment scoring is as follows: Alveolitis severity, bronchitis/bronchiolitis severity, rhinitis severity: 0 = no inflammatory cells, 1 = few inflammatory cells, 2 = moderate number of inflammatory cells, 3 = many inflammatory cells. Alveolitis extent, 0 = 0%, 1 = <25%, 2 = 25-50%, 3 = >50%. Alveolar oedema presence, alveolar haemorrhage presence, type II pneumocyte hyperplasia presence, 0 = no, 1 = yes. Extent of peribronchial/perivascular cuffing, 0 = none, 1 = 1-2 cells thick, 2 = 3-10 cells thick, 3 = >10 cells thick. Histopathological analysis was performed by Peter Wohlsein Dipl ECVP (Division of Pathology, University of Veterinary Medicine Hannover, Germany).

#### Statistical analysis

One-way ANOVA followed by Dunnett’s multiple comparisons tests were performed using GraphPad Prism version 9.0.0 for Windows, GraphPad Software, San Diego, California USA, www.graphpad.com

## Supplementary tables

**Table S1.** Summary of the quantification of viral load in throat swabs, lung and nasal turbinate tissue. Values are presented as Mean (SD).

| Group | Throat Swab titration (Log10 TCID<sub>50</sub>/ml) |  |  |  |  | Tissue titration (Log10 TCID<sub>50</sub>/g) |  |
| --- | --- | --- | --- | --- | --- | --- | --- |
|  | Day 0 | Day 1 | Day 2 | Day 3 | Day 4 | /g Lung | /g Turbinate |
| Vehicle | 0.80 (0) | 2.73 (0.45) | 3.55 (0.64) | 2.82 (0.37) | 1.75 (0.52) | 7.12 (0.39) | 7.98 (0.44) |
| 250mg/kg Epeleuton | 0.80 (0) | 1.08 (0.45) | 1.70 (1.01) | 2.23 (0.73) | 2.62 (0.70) | 5.78 (2.38) | 6.85 (2.32) |
| 500mg/kg Epeleuton | 0.80 (0) | 0.80 (0) | 0.80 (0) | 1.62 (1.13) | 2.98 (1.10) | 3.97 (2.39) | 6.32 (2.16) |

**Table S2.** Summary of viral RNA detected in throat, lung and nasal turbinate tissue. Values are presented as Mean (SD).

| Group | Throat Swab PCR (Log10 CP/ml) |  |  |  |  | Tissue PCR (Log10 CP/g) |  |
| --- | --- | --- | --- | --- | --- | --- | --- |
|  | Day 0 | Day 1 | Day 2 | Day 3 | Day 4 | /g Lung | /g Turbinate |
| Vehicle | 1.9 (0) | 5.58 (0.34) | 6.78 (0.57) | 5.88 (0.45) | 5.33 (0.56) | 10.52 (0.40) | 11.83 (0.22) |
| 250mg/kg Epeleuton | 1.9 (0) | 3.38 (0.88) | 4.18 (1.25) | 5.17 (0.55) | 5.55 (0.50) | 9.37 (1.71) | 10.68 (1.50) |
| 500mg/kg Epeleuton | 1.90 (0) | 2.45 (0.44) | 2.58 (0.58) | 3.90 (1.56) | 5.53 (1.32) | 7.60 (2.32) | 9.63 (2.56) |

**Table S3.** Scores of all assessed histopathological parameters.

| Group | n | Average Score |  |  |  |  |  |  |
| --- | --- | --- | --- | --- | --- | --- | --- | --- |
|  |  | Extent of alveolitis (0-3) | Severity of alveolitis (0-3) | Alveolar haemorrhage (% positive slides) | Severity of bronchi(ol)itis (score 0-3) | Degree of peribronchial/perivascular cuffing (score 0-3) | Severity of rhinitis (score 0-3) | Severity of tracheitis (score 0-3) |
| *Vehicle* | 6 | 0.83 | 1.33 | 17 | 2.17 | 1.67 | 2.83 | 2.33 |
| *Epeleuton 250mg/kg* | 6 | 0.33 | 0.50 | 17 | 0.83 | 0.50 | 2.00 | 2.00 |
| *Epeleuton 500mg/kg* | 6 | 0.17 | 0.17 | 0 | 0.33 | 0.17 | 1.17 | 1.00 |

